# Targeting ischemia-induced KCC2 hypofunction rescues refractory neonatal seizures and mitigates epileptogenesis in a mouse model

**DOI:** 10.1101/2020.09.15.298596

**Authors:** Brennan J. Sullivan, Pavel A. Kipnis, Brandon M. Carter, Shilpa D. Kadam

**Author notes:** **Corresponding Author**: Shilpa D. Kadam, PhD, Neuroscience Laboratory, Hugo Moser Research Institute at Kennedy Krieger; Department of Neurology, Johns Hopkins University School of Medicine, 707 North Broadway, 400H; Baltimore, MD 21205.

## Abstract

Neonatal seizures pose a clinical challenge for their early detection, acute management, and mitigation of long-term comorbidities. A major cause of neonatal seizures is hypoxic-ischemic encephalopathy that results in seizures that are frequently refractory to the first-line anti-seizure medication phenobarbital (PB). One proposed mechanism for PB-inefficacy during neonatal seizures is the reduced expression and function of the neuron-specific K^+^/Cl^−^ cotransporter 2 (KCC2), the main neuronal Cl^−^ extruder that maintains chloride homeostasis and influences the efficacy of GABAergic inhibition. To determine if PB-refractoriness after ischemic neonatal seizures is dependent upon KCC2 hypofunction and can be rescued by KCC2 functional enhancement, we investigated the recently developed KCC2 functional enhancer CLP290 in a CD-1 mouse model of refractory ischemic neonatal seizures quantified with vEEG. We report that acute CLP290 intervention can rescue PB-resistance, KCC2 expression, and the development of epileptogenesis after ischemic neonatal seizures. KCC2 phosphorylation sites have a strong influence over KCC2 activity and seizure susceptibility in adult experimental epilepsy models. Therefore, we investigated seizure susceptibility in two different knock-in mice in which either phosphorylation of S940 or T906/T1007 was prevented. We report that KCC2 phosphorylation regulates both neonatal seizure susceptibility and CLP290-mediated KCC2 functional enhancement. Our results validate KCC2 as a clinically relevant target for refractory neonatal seizures and provide insights for future KCC2 drug development.

## Introduction

Neonatal seizures occur in an estimated 1 to 3.5 per 1000 live births in the term infant. Hypoxic-ischemic encephalopathy (HIE) is a major cause of acute neonatal seizures (*1*). The management of these seizures are a major clinical challenge as they are often refractory to an initial loading dose of the first-line anti-seizure medication (ASM) phenobarbital (PB) and adjunct ASMs (*2, 3*). Compared to seizures at older ages, neonatal seizures differ in their etiology, semiology, electrographic signature, and response to ASMs (*1, 4*). There is an urgent need to identify the developmental mechanisms underlying seizure susceptibility and ineffective ASM response in the neonatal brain.

The neonatal brain has lower expression of its chief chloride extruder the neuron-specific K^+^-Cl^−^ cotransporter 2 (KCC2), and a high neuronal intracellular chloride concentration ([Cl^−^]_i_) (*5*–*7*). GABA is the primary inhibitory neurotransmitter in the mature brain, however in the neonatal brain the activation of GABA_A_ receptors (GABARs) results in depolarizing actions on immature neurons (*8*). KCC2 expression and function increase during development, resulting in a lower [Cl^−^]_i_ that coincides with a developmental shift from depolarizing to hyperpolarizing GABAergic signaling. The importance of KCC2 function in seizure susceptibility is supported by emerging evidence from human genetics, as pathogenic variants in *SLC12A5* are associated with the development of idiopathic generalized epilepsy and early infantile epileptic encephalopathy (OMIM #616685 and #616645, respectively) (*9*).

Despite the introduction of many new ASMs into clinical practice over the past 20 years, the incidence of refractory seizures has remained unchanged (*10*). KCC2 hypofunction is increasingly associated with pharmaco-resistant epilepsies and is a proposed cause of disinhibition (*11, 12*). GABA_A_R mediated fast synaptic inhibition and the anti-seizure efficacy of PB (a positive allosteric modulator of GABA_A_Rs) are both dependent upon active neuronal Cl^−^ extrusion (*13, 14*). In a preclinical CD-1 mouse model of ischemic neonatal seizures, KCC2 is transiently downregulated and associated with PB-resistant seizures (*15*). In this model, post-ischemic tropomyosin receptor kinase (TrkB) inhibition rescued PB-resistant neonatal seizures and KCC2 expression (*16, 17*). This acute rescue of KCC2 hypofunction via TrkB inhibition improved long-term outcomes after ischemic neonatal seizures (*18, 19*). These results suggest KCC2 as a novel therapeutic target for refractory neonatal seizures.

A recent high throughput screen for compounds that reduce [Cl^−^]_i_ in a cell line with low KCC2 expression identified the compound CLP257 as a KCC2 functional enhancer (*20*). To improve the poor pharmacokinetics of CLP257, the carbamate prodrug CLP290 was designed and improved the half-life (t_1/2_) from <15min to 5h (*20*). These compounds provide an opportunity to test whether KCC2 functional enhancement can rescue the emergence of PB-resistant neonatal seizures after ischemia in our preclinical CD-1 mouse model. KCC2 expression increases twofold in mouse pups during development between P7 and P10 (*15*).Therefore to examine the developmental differences in ischemic seizure suppression after KCC2 functional enhancement both age groups were included. Neonatal seizures are associated with poor long-term outcomes including the development of epilepsy (*21, 22*). Therefore, the effect of acute CLP290 intervention at P7 was evaluated at P12 using a new subthreshold pentylenetetrazol (PTZ) dosing protocol to investigate epileptogenesis. To test if KCC2 hypofunction could induce ictal events independent of ischemia, the selective KCC2 inhibitor VU0463271 was administered to P7 naïve pups during vEEG. KCC2 posttranslational modifications are tightly regulated throughout development and strongly influence KCC2 activity (*23*–*25*). The KCC2 phosphorylation sites S940 (*25*) and T1007 (*24*) were investigated for their role in CLP290-mediated effects on neonatal refractory seizures. Using two knock-in mutant mice that prevent either S940 (*25*) or T1007 (*24*) phosphorylation, we investigated the importance of these sites on neonatal seizure susceptibility and post-ischemic PB-efficacy.

## Results

### CLP290 rescues phenobarbital-resistant seizures at P7

The true burden of acute ischemic seizures in mouse pups cannot be identified by behavioral scoring parameters alone and require continuous vEEG recordings as their presentation can range from entirely electrographic to generalized convulsive (*7*). In our CD-1 neonatal mouse model of unilateral carotid ligation, pups presented with PB-resistant ischemic neonatal seizures at P7 (Fig. 1A-D), an established characteristic of the model (*15*–*17, 26*). All pups underwent 1h of baseline vEEG recording after unilateral carotid ligation. After 1h of vEEG recording all pups received a loading dose of PB with a subsequent hour of vEEG recording (Fig. 1A-D). Pups that only received a PB loading dose (PB-only) were pharmacoresistant, as their 2^nd^h seizure burden remained high and was similar to 1^st^h levels (Fig 1C-F).

**Fig. 1.**
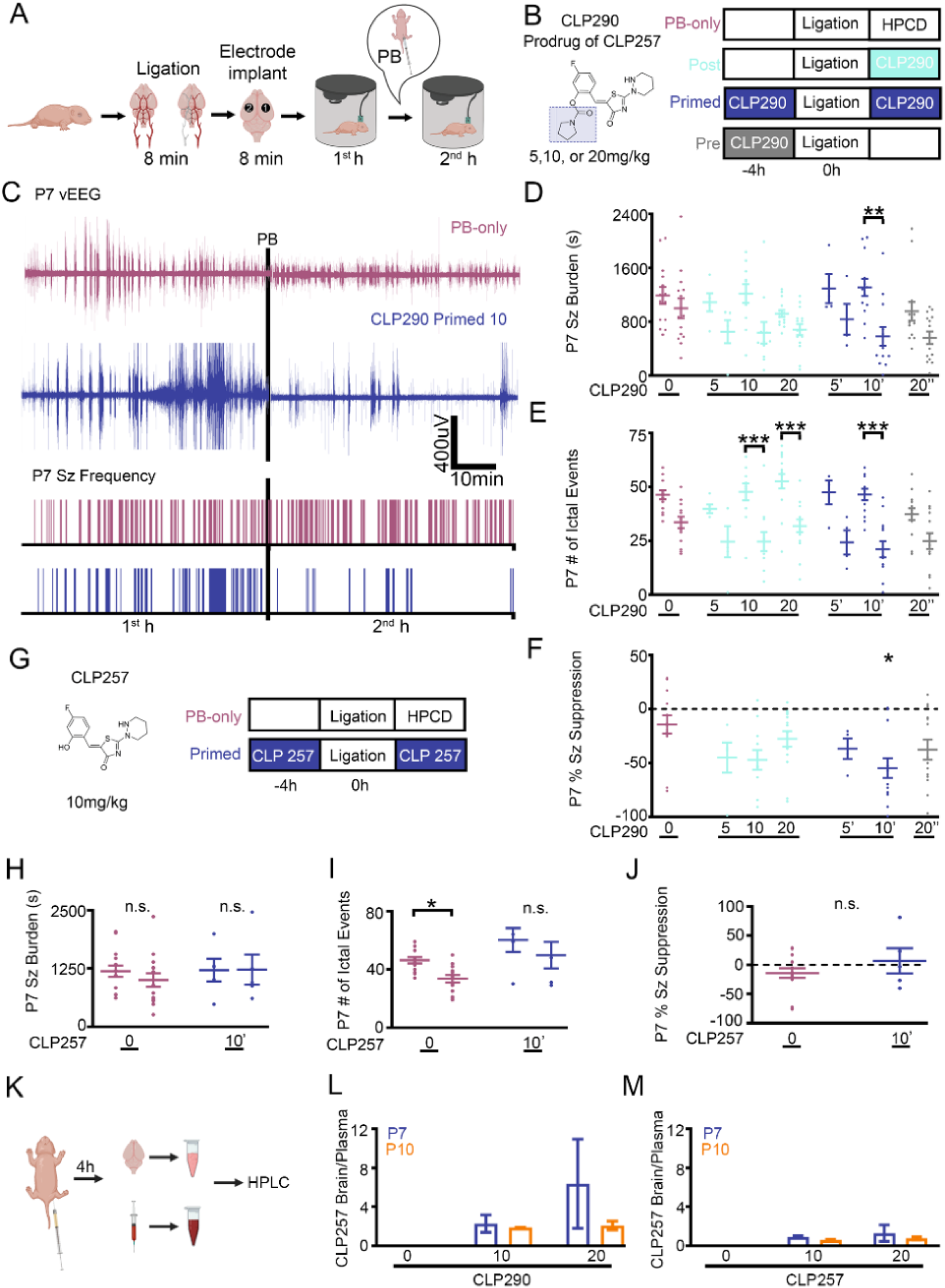
CLP290 rescued phenobarbital-resistant neonatal seizures in P7-CD1 mice. (**A**) Experimental design of a P7 CD-1 mouse model of ischemic neonatal seizures with continuous vEEG. Recording (1) and reference (2) electrodes over bilateral parietal cortices, with a ground electrode over the rostrum. (**B**) Doses and treatment protocols to evaluate CLP290, a prodrug of the proposed KCC2 functional enhancer CLP257. (**C**) Representative EEG traces and seizure frequency raster plots of a PB-only and CLP290 10’ pup. Black bars represent a loading dose of PB (25mg/kg; intraperitoneal injection). (**D**) 1^st^ and 2^nd^ hour seizure burdens, **(E)** 1^st^ and 2^nd^ hour ictal events, and (**F**) 1^st^ vs. 2^nd^ hour percent seizure suppression after P7 unilateral carotid ligation. Percent seizure suppression was analyzed by one-way ANOVA vs. PB-only. PB-only n=14; CLP290 5 n=5; CLP290 10 n=11; CLP290 20 n=15; CLP290 5’ n=4; CLP290 10’ n=13; CLP290 20’ n=14. (**G**) Doses and treatment protocols to evaluate CLP257 (n=5). (**H**) Seizure burdens, (**I**) ictal events, and (**J**) percent seizure suppression at P7 after unilateral carotid ligation. (**K**) Experimental paradigm to investigate the pharmacokinetic profile of CLP290 and CLP257. (**L**) CLP257 brain to plasma ratio after CLP290 administration (I.P., n=2 per group). (**M**) CLP257 brain to plasma ratio after CLP257 administration (I.P.; n=2 per group). Plots show all data points with means ±SEM. *P<0.05; **P<0.01; ***P<0.001, two-way ANOVA.

To investigate if the KCC2 functional enhancer CLP290 could rescue PB-resistant seizures, CLP290 was administered intraperitoneally at P7 in the Pre, Post, or Primed groups (Fig. 1 B). Administration of Primed CLP290 10mg/kg (10’) significantly reduced total seizure burden, duration, and frequency of 2^nd^h seizure events (Fig. 1C-E). This decrease in 2^nd^h seizure burden significantly increased seizure suppression after PB administration, thereby rescuing P7 PB-resistance (Fig. 1D-F). In contrast to 10’, bolus administration of 20mg/kg CLP290 reduced 1^st^h baseline seizure burden (Supplemental Fig. 1).

### Pro-drug CLP290 improved brain availability

A major limiting factor for epilepsy drug development is the *in vivo* brain availability of candidate compounds identified *in vitro* (*27, 28*). CLP290 is a carbamate prodrug of CLP257 and has an improved t_1/2_ from <15min (CLP257) to 5h (CLP290) in blood samples from adult rats (*20*). The brain availability of CLP257 and CLP290 have not been previously published. At P7, CLP257 was unable to reduce 2^nd^h seizure burden (Fig. 1 G-J) when administered systemically at the 10’ dose. To determine if CLP290 efficacy and CLP257 inefficacy were due to differences in pharmacokinetics, we performed HPLC to analyze brain levels of CLP257 after I.P. delivery of either CLP257 or CLP290 (Fig. 1K and Supplemental Fig. 2). At P7 and P10, CLP290 administration of 10 or 20mg/kg demonstrated adequate brain availability. (Fig. 1L). CLP257 had poor brain availability compared to CLP290 (Fig. 1K-M), which suggested that the inability of CLP257 to rescue PB-refractory seizures was due to its poor pharmacokinetic profile.

**Fig. 2.**
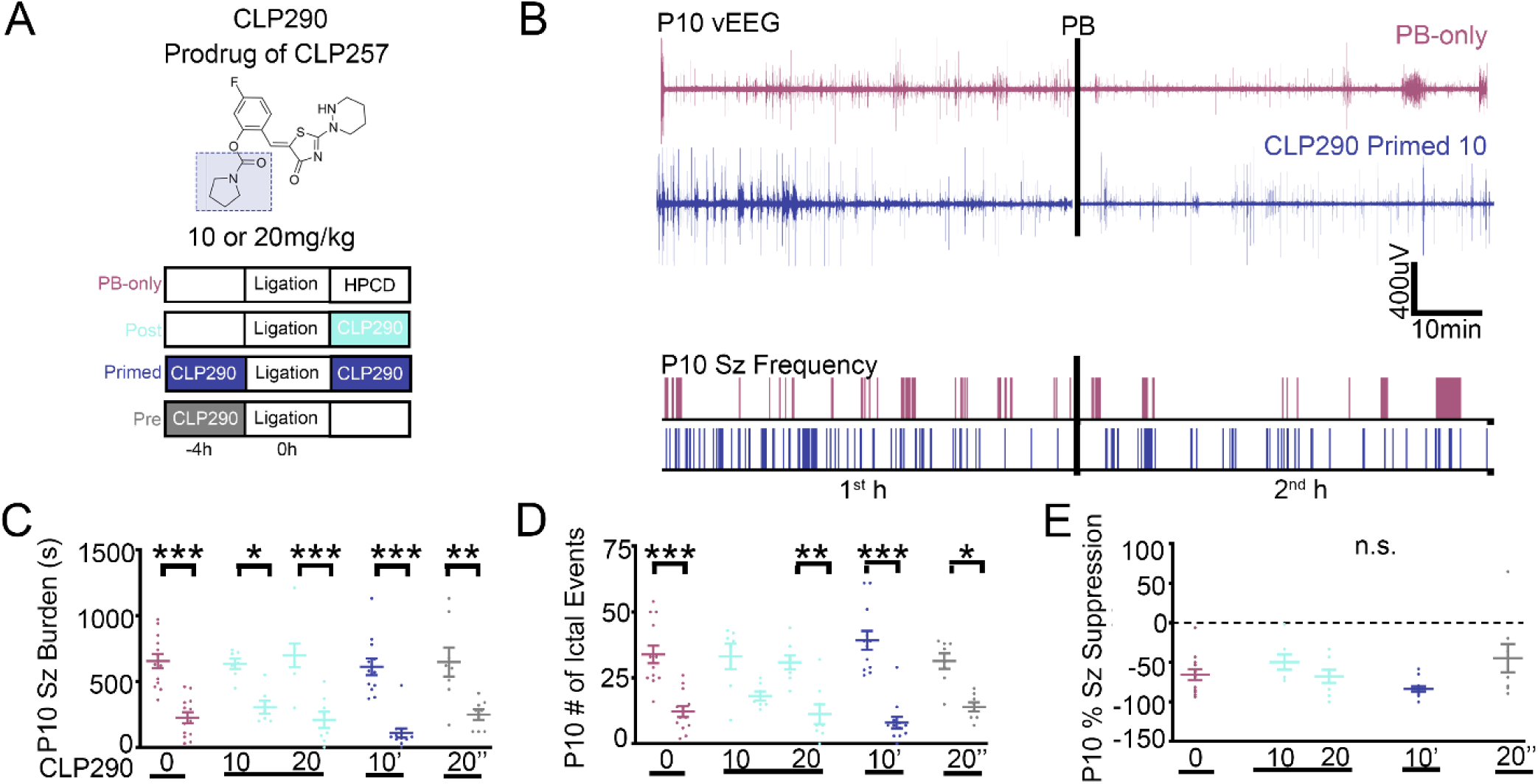
Ischemic neonatal seizures in P10_-CD1 pups were phenobarbital-responsive. (**A**) Doses and treatment protocols to evaluate CLP290 in P10 CD-1 pups. (**B**) Representative EEG traces and seizure frequency raster plots of a P10 phenobarbital-only and CLP290 10’ pup. Black bars represent a loading dose of PB (25mg/kg; intraperitoneal injection). (**C**) 1^st^ and 2^nd^ hour seizure burdens, **(D)** 1^st^ and 2^nd^ hour ictal events. (**E**) CLP290 does not improve seizure suppression at P10. 1^st^ vs. 2^nd^ hour percent seizure suppression after unilateral carotid ligation at P10. Percent seizures suppression was analyzed by one-way ANOVA vs. PB-only. Plots show all data points with means ±SEM. *P<0.05; **P<0.01; ***P<0.001, two-way ANOVA. PB-only; n=13; CLP290 10 n=7; CLP290 20 n=8; CLP290 10’ n=12; CLP290 20” n=8.

### CLP290 efficacy on phenobarbital-responsive seizures at P10

As previously reported (*15*–*17, 26*), ischemic neonatal seizures after unilateral carotid ligation in P10 CD-1 pups were PB-responsive (Fig. 2 A-E). The age-dependent emergence of PB-resistant and PB-responsive seizures at P7 versus P10 is a characteristic of the neonatal mouse model that is associated with a twofold increase in KCC2 expression between P7 and P10 (*15*). Administration of CLP290 at P10 did not further improve the efficacy of PB at any of the doses tested (Fig. 2 C-E). When ischemic neonatal seizures were PB-responsive at P10, the previously reported TrkB-antagonist ANA12 (*17, 26*) also did not improve the efficacy of PB. This suggests that the therapeutic benefit of KCC2 functional enhancement is dependent upon the degree of KCC2 hypofunction which is evident in the emergence of refractoriness at P7 but not at P10.

### CLP290 rescued ipsilateral post-ischemic KCC2 and S940 downregulation

PB-resistant ischemic neonatal seizures have been shown to significantly reduce expression of KCC2 and phosphorylation of S940 24h after ischemia (*17*). P7 unilateral carotid ligation did not result in an infarct stroke injury, therefore KCC2 degradation at 24h is not caused by infarct related cell-death (*15*). In this model, KCC2 expression undergoes a recovery over 3-4 days, a characteristic that is associated with the transient nature of neonatal seizures due to HIE (*29, 30*). At 24h after P7 unilateral carotid ligation, KCC2 and S940 expression was significantly lower in the right hemisphere (ipsilateral to ischemia) than the left hemisphere (contralateral to ischemia) in the PB-only group (Fig. 3 A-E). Intervention with CLP290 at P7 rescued both KCC2 and S940 downregulation in the 10’ group 24h after ligation (Fig. 3 D-E). Bolus administration of CLP290 20mg/kg Post significantly increased S940 expression bilaterally compared to the PB-only group at P7 (Fig. 3C and Supplemental Fig. 3 A-B). When ischemic neonatal seizures are PB-responsive at P10, KCC2 expression was not significantly decreased at 24h (Fig. 3 F-I). However, the ratio of ipsilateral to contralateral S940 expression in the PB-only group was significantly decreased (Fig. 3 H-J) and was rescued in all CLP290 treatment groups (Fig. 3 H-J). Therefore, when ischemic neonatal seizures were PB-resistant, CLP290 rescued KCC2 expression and S940 phosphorylation, suggesting that the degree of KCC2 hypofunction drives the therapeutic benefit of KCC2 functional enhancement.

**Fig. 3.**
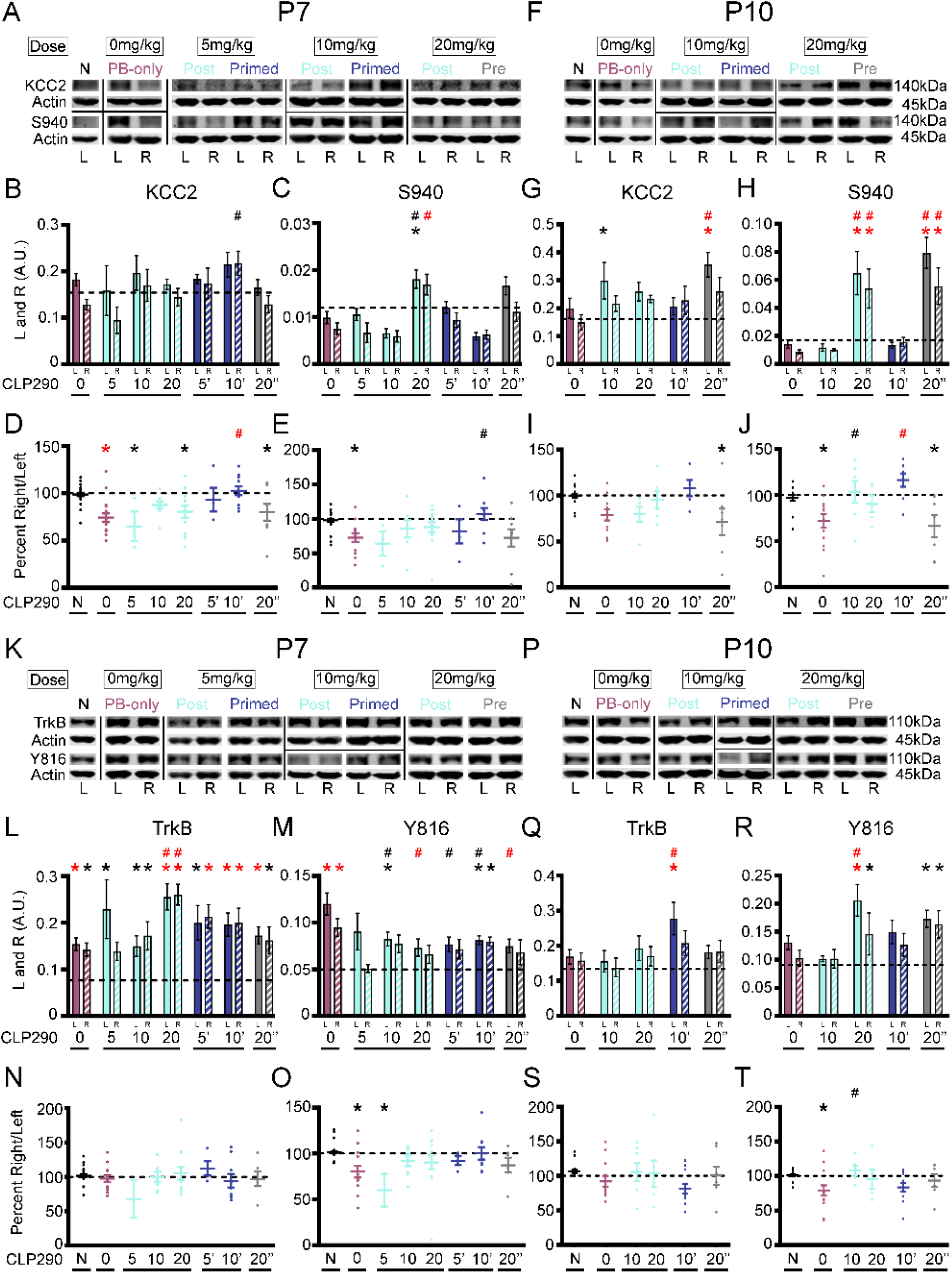
CLP290 rescued post-ischemic KCC2 downregulation but not TrkB activation. (**A**) Representative Western blots of KCC2 and S940 protein expression 24h after P7 ischemic seizures. (**B**) KCC2 and (**C**) S940 expression in left (L) and right (R) hemispheres. (**D**) KCC2 and (**E**) S940 expression as percent ipsilateral/contralateral (R/L). (**F**) Representative Western blots of KCC2 and S940 expression 24h after P10 ischemic seizures. (**G**) KCC2 and (**H**) S940 expression in L and R hemispheres. (**I**) KCC2 and (**J**) S940 expression as percent R/L. (**K**) Representative Western blots of TrkB and Y816 expression 24h after P7 ischemic seizures. (**L**) TrkB and (**M**) Y816 expression in L and R hemispheres. (**N**) TrkB and (**O**) Y816 expression as percent R/L. (**P**) Representative Western blots of TrkB and Y816 expression 24h after P10 ischemic seizures. (**Q**) TrkB and (**R**) Y816 expression in L and R hemispheres. (**S**) TrkB and (**T**) Y816 expression as percent R/L. Data plots show means ±SEM. All proteins of interest were normalized to housekeeping protein β-actin. Phosphoproteins were normalized to their respective total protein. *P<0.05 and *P<0.001 by 1-way ANOVA vs. Naive. #P<0.05 and #P<0.001 vs. PB-Only. P7 pups: Naïve n=27; PB-only n=18; 5 Post n=3; 10 Post n=9; 20 Post n=13; 5’, n=4; 10’, n=11; 20 Pre n=9. P10 pups: Naïve n=18, PB-only n=11, 10 Post n=6, 20 Post n=6, 10 Primed n=5, 20 Pre n=7.

### CLP290 rescue of PB-resistant seizures was not mediated through BDNF-TrkB

Ischemic neonatal seizures at P7 induce a significant bilateral increase in TrkB expression and phosphorylation of Y816 (*17, 26*). Similarly, TrkB expression and Y816 phosphorylation were significantly upregulated bilaterally in the PB-only group 24h after ischemic neonatal seizures at P7 (Fig. 3K-O) but not at P10 (Fig. 3 P-T and Supplemental Fig. 3 C-D). In this study, the bilateral increase in post-ischemic TrkB and Y816 expression was detected as previously characterized (*17*). Increased TrkB expression was not significantly rescued by CLP290 however Y816 was (Fig. 3 K-O). 24h after P7, the Y816/TrkB ratios demonstrated a significant bilateral reduction in the ratio by CLP290 20 post (Supplemental Fig. 3C), driven by the significant increase in TrkB expression (Fig. 3L-O). In this model, ischemic neonatal seizures at P10 were PB-responsive and P10 pups did not show activation of TrkB (Fig. 3P-T and Supplemental Fig. 3 C-D). These results suggest that the CLP290-mediated seizure suppression in the 10’ group at P7 was independent of TrkB.

### Acute CLP290 intervention at P7 mitigates epileptogenesis at P12

HIE seizures are transient within the first week of life (*29, 30*) and the long-term consequences of neonatal seizures are difficult to isolate from the consequences of prolonged and/or inefficacious anti-seizure therapy in the clinic (*22*). In our CD-1 mouse model of refractory neonatal ischemic seizures, we have previously documented the emergence of epilepsy in adulthood (*31*). To assess if the acute rescue of PB-resistant neonatal seizures at P7 via CLP290 had long-term benefits, the same pups underwent a PTZ challenge at P12 (Fig. 4A-B). Previously, a 80mg/kg dose of PTZ was shown to induce high seizure burdens that were PB-responsive and upregulated KCC2 at P7 in CD-1 pups (*32*). In this study, pups were administered three doses of PTZ (20, 20, and 40 mg/kg) 1h apart (Fig. 4A-B). Using this protocol, we characterized seizure susceptibility in P7 naïve, P7 PB-only, and P7 CLP290 10’ treated pups at P12. The initial dose of PTZ induced seizures in the 1^st^h for all groups (Fig. 4A-D). Naïve pups demonstrated a general reduction in seizures during the second and third PTZ doses, potentially uncovering a homeostatic compensation to help develop resistance to the chemoconvulsant induced seizures. PB-only pups demonstrated an increase in seizure burden after the repeated doses of PTZ, with mice (n=3) going into status epilepticus. Thus, the significant increase in PB-only seizure burdens at P12 was driven by a significant increase in seizure duration (Fig. 4E). CLP290 10’ treated pups had similar P12 seizure burdens as naïve pups and significantly lower than PB-only pups (Fig. 4A-D). This novel PTZ challenge protocol identified epileptogenesis in the form of heightened susceptibility to seizures at P12 for neonatal pups that underwent standard but inefficacious PB treatment for their refractory seizures, efficacious CLP290 10’ intervention at P7 resulted in the regression of epileptogenesis at P12.

**Fig. 4.**
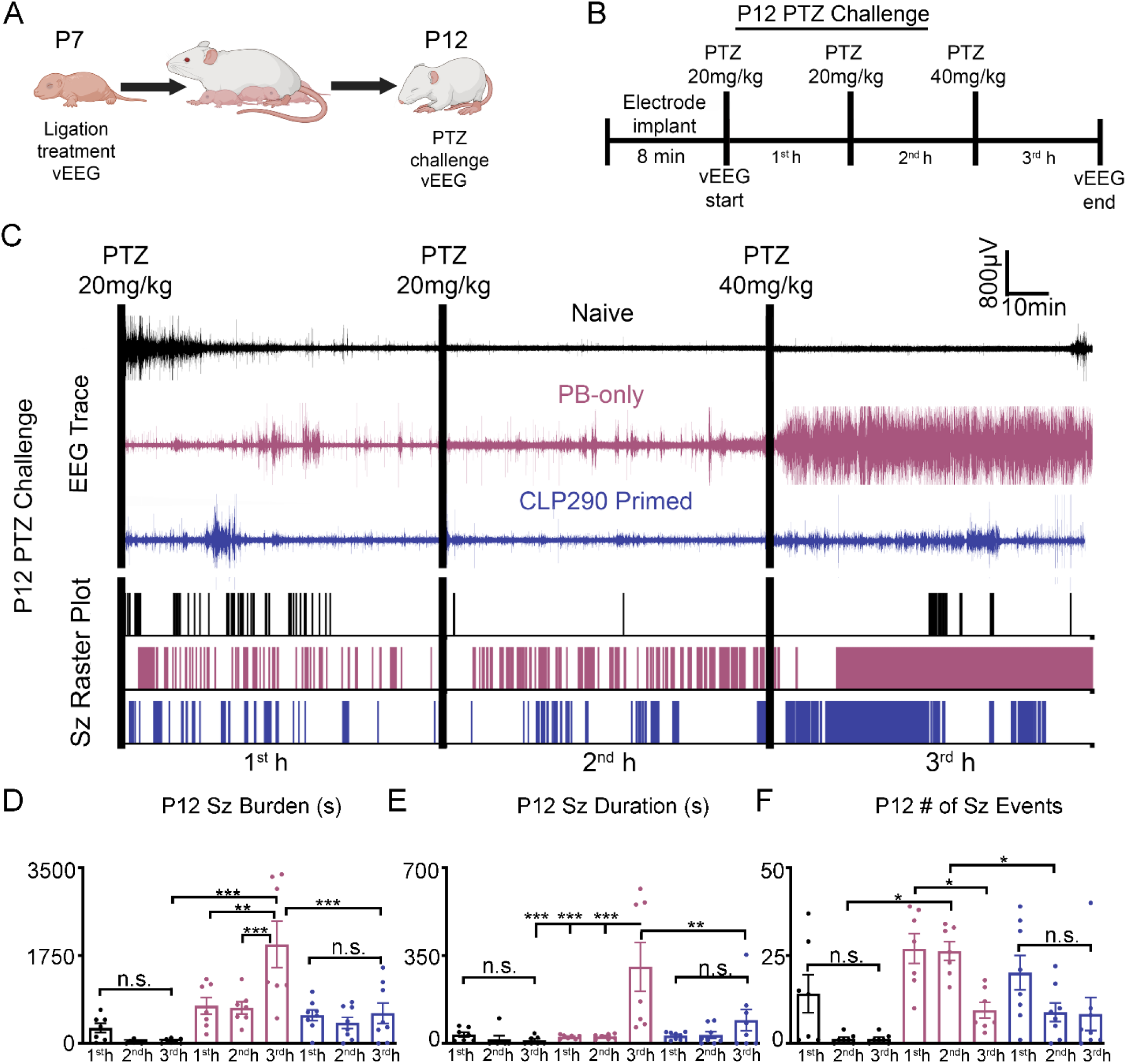
CLP290-mediated regression of epileptogenesis detected using a PTZ challenge. (**A**) Schematic to investigate the developmental benefits of CLP290 10’ treatment for P7 ischemic seizures. (**B**) P12 pentylenetetrazol (PTZ) challenge to evaluate epileptogenesis. (**C**) Representative EEG traces and seizure frequency raster plots for P12 pups that underwent Naïve, PB-only, or CLP290 10’ treatment at P7. Black bars indicate intraperitoneal PTZ injections. (**D**) 1^st^, 2^nd^, and 3^rd^ hour seizure burdens at P12 in CD-1 mice after PTZ injections. (**E**) Total electrographic seizure burdens over the three hours of vEEG recording. (**F**) Total seizure events over three hours of vEEG recording. Data plots show all data points with means ±SEM. *P<0.05; **P<0.01; ***P<0.001 by 2-way ANOVA. Naïve n=7; PB-only n=7; CLP290 10’ n=8.

### In vivo KCC2 inhibition is epileptogenic in the neonatal brain

The selective KCC2 inhibitor VU0463271 (VU) has been shown to induce epileptiform discharges in the dorsal hippocampus of the adult mouse, highlighting the critical role of KCC2 in the mature hippocampus (*33*). Here, naive P7 CD-1 pups were administered 0.25mg/kg VU (I.P.) at the initiation of vEEG recording with a subsequent dose of 0.5mg/kg VU (I.P.) at 1h (Fig. 5A-B). Selective inhibition of KCC2 was sufficient to induce epileptiform activity (Fig. 5A-E) at P7. Prolonged and repeated seizures are known to play a role in the reduction of neuronal surface KCC2 expression and function (*15, 34*–*36*). At P7, if post-ischemic KCC2 hypofunction plays a critical role in PB-refractoriness, KCC2 inhibition following repeated ischemic seizures would be expected to further aggravate the seizure burden. To test the effect of KCC2 inhibition following repeated ischemia-induced neonatal seizures at P7, VU 0.25 mg/kg (I.P.) was administered 1h after unilateral carotid ligation (Fig. 5F). P7 pups that received VU after 1h of ischemic seizures developed a significant aggravation of EEG seizure burden in the second hour (Fig. 5F-J). Taken together, KCC2 inhibition induced epileptiform activity in the naïve neonatal brain and exacerbated ischemic neonatal seizures at P7.

**Fig. 5.**
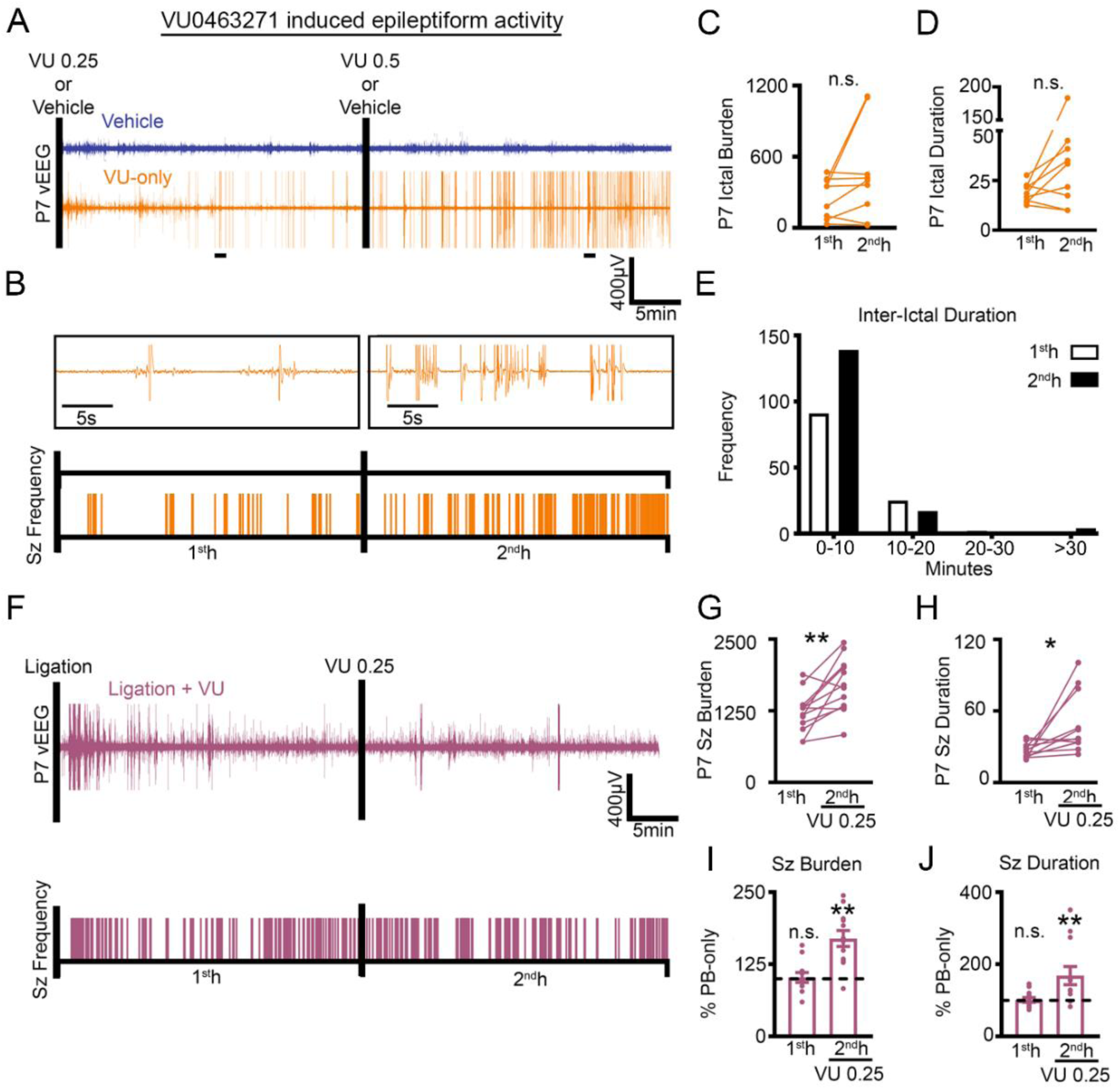
Selective KCC2 antagonist VU induced spontaneous epileptiform discharges in P7 pups. (**A**) Representative EEG traces and (**B**) ictal event frequency raster plots from P7 CD-1 pups that either underwent vehicle or VU0463271 (VU) administration. Black bars represent intraperitoneal injections. Expanded timescales show VU induced epileptiform activity in the first and second hour. (**C**) 1^st^ vs. 2^nd^ hour ictal burden and (**D**) ictal duration after VU administration. (**E**) Total frequency distribution for all interictal durations in P7 CD-1 pups administered VU. Vehicle (n=4) VU administration (n=8). (**F-J**) VU aggravated ischemic neonatal seizure burdens. (**F**) Representative EEG trace and seizure frequency raster plot of a P7 CD-1 pup that underwent unilateral carotid ligation with administration of VU 0.25mg/kg at 1h (denoted by black bar). (**G**) 1^st^ and 2^nd^ h seizure burden and (**H**) 1^st^ and 2^nd^ h seizure duration for P7 CD-1 pups that underwent unilateral carotid ligation with administration of VU0.25mg/kg at 1h. (**I**) 1^st^ and 2^nd^ hour seizure burden and (**J**) duration plotted as percent PB-only. Data plots show all data points with means ±SEM. **P<0.05 and **P<0.01 by two-tailed paired t-test. Ligation + VU n=12.

### In vitro CLP257 incubation increases KCC2 membrane insertion

The KCC2 functional enhancer CLP257 has been shown to increase chloride extrusion capacity and KCC2 membrane expression *in vitro*, and it has been suggested that the net effect of KCC2 functional enhancement may emerge from relatively small changes in KCC2 function (*11, 20, 37*). The neonatal brain tightly regulates KCC2 activity via S940 and T1007 phosphorylation (*23*), and it is unknown if age-dependent mechanisms affect KCC2 functional enhancement. Therefore, P7 neonatal brain sections were incubated with graded doses of CLP257 or CLP290 (Fig. 6A-C). 500μM CLP257 significantly increased membrane KCC2 and S940 expression (Fig. 6B). The phosphorylation of S940 increases KCC2 plasma membrane accumulation and transport activity (*38, 39*). To assess if the S940 site was necessary for CLP257 mediated KCC2 membrane insertion, brain sections from S940A^+/+^ knock-in mutant mice (*25*) were treated with CLP257 (Fig. 6 D-G). Without the ability to phosphorylate S940, CLP257 failed to increase KCC2 membrane insertion (Fig. 6 F).

**Fig. 6.**
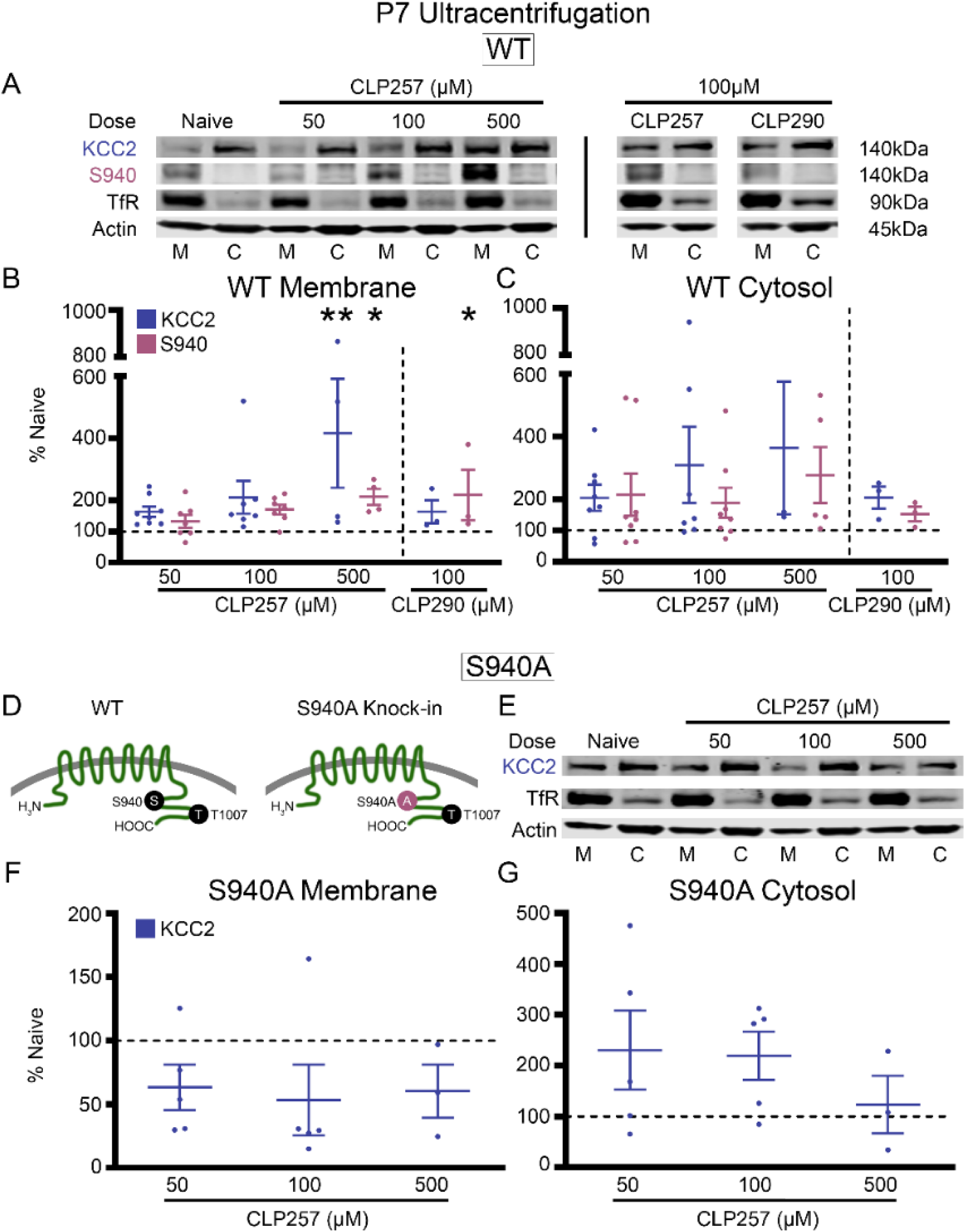
CLP257 upregulated membrane KCC2 expression and S940 phosphorylation. (**A**) KCC2 and S940 protein expression in the plasma membrane (M) and cytosol (C) for all treated P7 wildtype (WT) brain slices. (**B**) KCC2 and S940 protein expression in the membrane and (**C**) cytosol for all treatment groups plotted as percent of naïve. Number of WT P7 pups: n=8 (50μM CLP257), n=7 (100μM CLP257), n=4 (500μM CLP257), n=3 (100μM CLP290). KCC2 functional enhancement by CLP257 is dependent upon the phosphorylation of S940. (**D**) Graphical representation of S940A^+/+^ knock-in mutant mice (*36*). (**E**) KCC2 expression in the plasma membrane and cytosol for all treated brain slices from S940A^+/+^ P7 pups. (**F**) KCC2 expression in the membrane and (**G**) cytosol for all S940A^+/+^ treatment groups plotted as percent of naïve. All proteins of interest in the cytosol were normalized to housekeeping protein β-actin. All proteins of interest in the plasma membrane were normalized to transferrin (TfR). Phosphoproteins were normalized to their respective total protein. Data plots show all data points with means ±SEM. * P<0.05 and ** P<0.01 by 1-way ANOVA. S940A^+/+^ P7 pups: n=5 (50μM CLP257), n=5 (100μM CLP257), n=3 (500μM CLP257).

### Ischemic neonatal seizures do not modulate KCC2-T1007 phosphorylation

The phosphorylation of KCC2 residue T1007 inhibits KCC2 function (*40, 41*). It has been proposed that KCC2 functional enhancers must either increase KCC2 surface stability and/or decrease T1007 phosphorylation (*42, 43*). Therefore, T1007 phosphorylation was investigated 24h after P7 ischemic neonatal seizures (Fig. 7A-C). Ischemia did not significantly change T1007 expression at 24h in either hemisphere (Fig. 7A-C and Supplemental Fig. 4). The effective CLP290 10’ dose also did not significantly modulate T1007 expression, however bolus administration of 20 mg/kg CLP290 resulted in a significant increase in T1007 at 24h in both pre and post treatment groups (Fig. 7A-D). To investigate if pharmacomodulation of KCC2 is governed by T1007, naïve CD-1 P7 brain sections were treated with CLP257 (Fig. 7E-H). All groups treated with CLP257 showed lower T1007 expression at the membrane than untreated slices (Fig. 7G) supporting the data from the *in vivo* experiments (Fig. 7B &C).

**Fig. 7.**
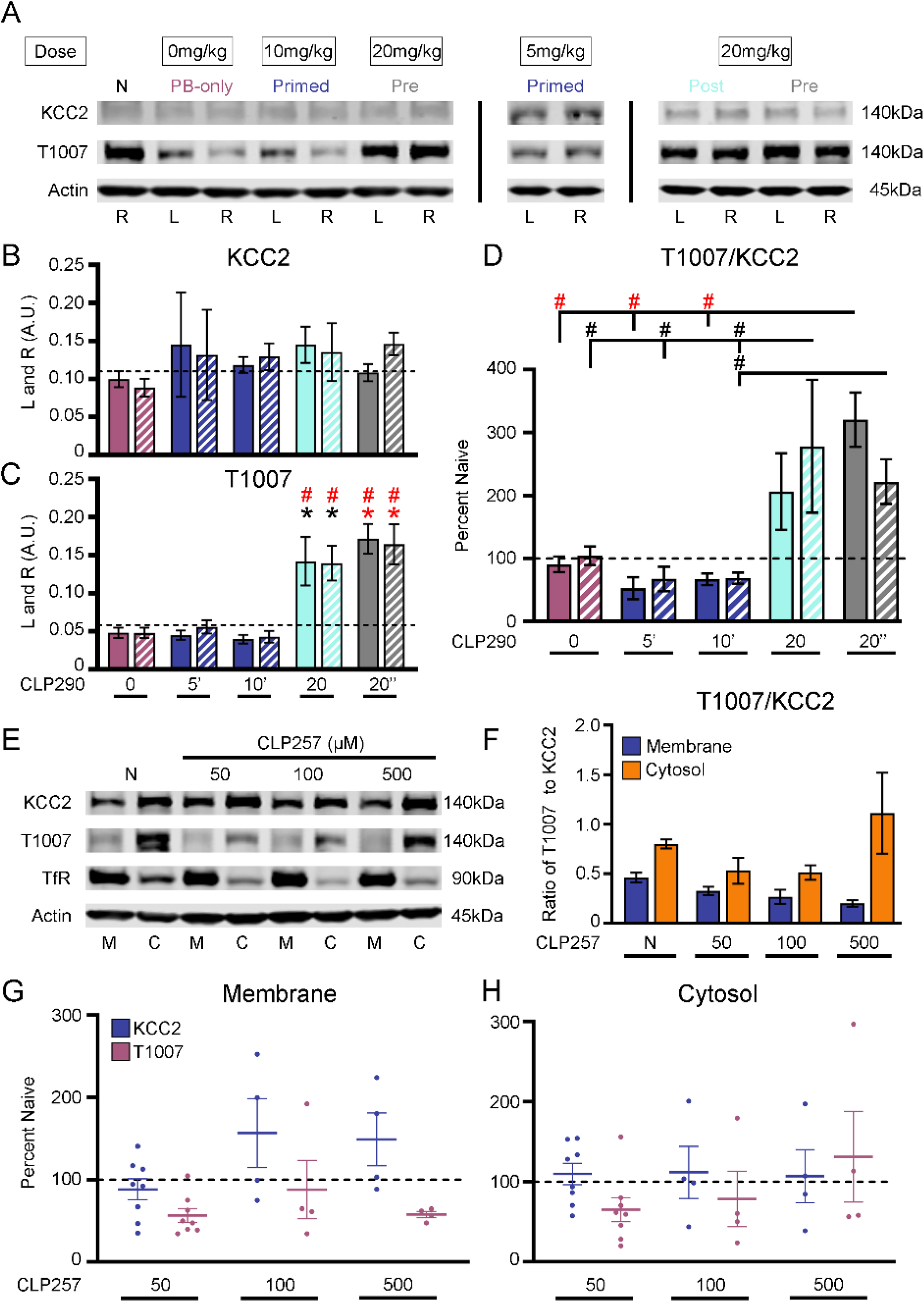
Bolus administration of CLP290 20mg/kg induced homeostatic upregulation of T1007 phosphorylation. (**A**) Representative western blot of KCC2 and T1007 expression 24h after ischemic neonatal seizures. (**B**) KCC2 and (**C**) T1007 expression in left (L) and right (R) hemispheres. (**D**) T1007/KCC2 ratios plotted as percent naïve for L and R hemispheres. Naïve n=10 PB-only; n=14; CLP290 5’ n=3; CLP290 10’ n=9; CLP290 20 n=4; CLP290 20” n=8. (**E**) Representative western blot of KCC2 and T1007 expression at the membrane (M) and cytosol (C) in CLP257 treated slices. (**F**) T1007-KCC2 ratio for M and C for CLP257 treated slices. (**G**) Membrane KCC2 and T1007 plotted as percent naïve. (**H**) Cytosol KCC2 and T1007 plotted as percent naïve. *P<0.05 and *P<0.001 by one-way ANOVA vs Naive. #P<0.05 and #P<0.001 by one-way ANOVA vs PB-Only. n=8 (50μM CLP257); n=4 (100μM CLP257); n=4 (500μM CLP257).

### Spontaneous epileptiform discharges in S940A^+/+^ pups

KCC2 deficient mice die postnatally with generalized seizures and respiratory failure (*44, 45*). S940A^+/+^ mice are susceptible to death after kainate induced status epilepticus in adulthood (*36*). In patients with idiopathic generalized epilepsy and early childhood onset of febrile seizures, heterozygous missense variants in *SLC12A5* have been identified (*46, 47*) and associated with a reduction in S940 phosphorylation (*46*). However, it is unknown if the prevention of S940 phosphorylation alone can induce spontaneous epileptiform activity during the neonatal period. Therefore, S940A^+/+^ pups underwent vEEG at P7 and P12 (Fig. 8A-D). S940A^+/+^ mice had spontaneous epileptiform discharges at both P7 and P12, which failed to respond to CLP290 intervention (Fig. 8AD). These results suggest that the prevention of S940 phosphorylation is sufficient for spontaneous epileptic activity during development.

**Fig. 8.**
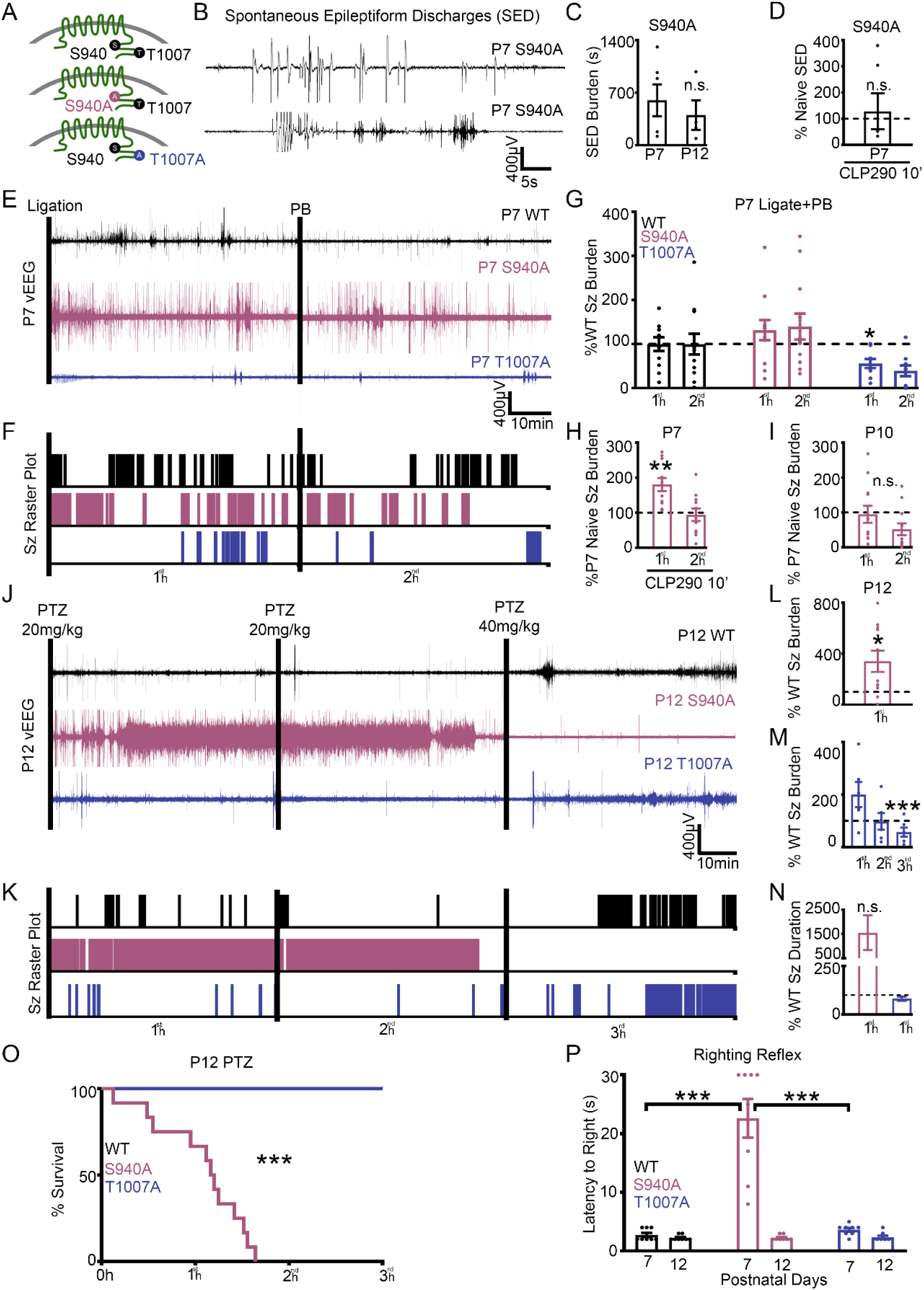
Inability to phosphorylate S940 or T1007 in knock-in pups regulated neonatal seizure susceptibility. (**A**) Graphical representation of homozygous S940A^+/+^ knock-in mutant mice (*25*) and homozygous T1007A knock-in mutant mice (*24*). (**B**) S940A^+/+^ mice had spontaneous epileptiform discharges (SEDs) at P7 and (**C**) P12. (**D**) Spontaneous epileptiform discharge duration of P7 S940A^+/+^ pups administered CLP290 10’ treatment. Naïve P7 S940A^+/+^ n=6; Naïve P12 S940A^+/+^ n=4; CLP290 10’ P7 S940A^+/+^ n=6. (E-G) P7 T1007A^+/+^ pups are resistant to ischemic neonatal seizures. (**E**) EEG traces and (**F**) seizure frequency raster plots for P7 WT, S940A^+/+^, and T1007A^+/+^pups that underwent unilateral carotid ligation. (**G**) 1^st^ vs 2^nd^ hour seizure burdens after unilateral carotid ligation plotted as percent WT. WT n=12; S940A^+/+^ n=12; and T1007A^+/+^ n=9. (**H**) P7 S940A^+/+^ ischemic seizures were CLP290 (10’) resistant. Seizure burden of CLP290 10’ treated P7 S940A^+/+^ pups after unilateral carotid ligation plotted as P7 percent naïve S940A^+/+^. CLP290 10’ treated P7 S940A^+/+^n=11. (**I**) Seizure burden of P10 S940A^+/+^ pups (n=8) after unilateral carotid ligation, plotted as P7 percent naïve S940A^+/+^. **(J-O)** Naïve P12 S940A^+/+^ mice are susceptible to status epilepticus and mortality. (**J**) Representative EEG trace and (**K**) seizure frequency raster plot of naïve P12 WT, S940A^+/+^, and T1007A^+/+^ pups. Black bars indicate intraperitoneal pentylenetetrazol (PTZ) injections. (**L**) Total seizure burden of P12 S940A^+/+^ pups after PTZ administration plotted as percent WT. (**M**) 1^st^, 2^nd^, and 3^rd^ hour seizure burdens at P12 in T1007A^+/+^ pups plotted as percent WT. (**N**) 1^st^ hour average seizure durations for S940A^+/+^ and T1007A^+/+^ P12 pups, plotted as percent WT. (**O**) Survival plot for P12 WT, T1007A^+/+^, and S940A^+/+^ pups during the PTZ challenge. P12 WT n=11; S940A^+/+^ n=11; and T1007A^+/+^ n=6 pups. (**P**) Duration of time to righting reflex at P7 and P12 in naïve WT, T1007A^+/+^, and S940A^+/+^ pups. (n=8 each group). *P<0.05; **P<0.01; ***P<0.001 by unpaired t-test vs. WT for all seizure data. Survival analysis ***P<0.001 by Mantel-Cox test. Righting Reflex by one-way ANOVA.

### T1007A^+/+^ pups are resistant to ischemic seizures

T1007 phosphorylation decreases during development and is associated with an increase in neuronal Cl^−^extrusion capacity (*23, 48, 49*). T1007A^+/+^ knock-in mutant mice have a reduced susceptibility to kainate induced status epilepticus in adulthood (*24*). It is unknown if KCC2 phosphorylation alone can modulate the susceptibility to ischemic neonatal seizures. To investigate the role of KCC2 phosphorylation in ischemic neonatal seizures WT, S940A^+/+^, and T1007A^+/+^ P7 pups underwent unilateral carotid ligation with 2h vEEG recordings (Fig. 8E-G). After P7 unilateral carotid ligation T1007A^+/+^pups were significantly resistant to ischemic seizures than WT (Fig. 8E-G). This result suggests that reducing T1007 phosphorylation on KCC2 is a promising therapeutic target to be developed for neonatal seizures. S940A^+/+^ pups had higher seizure burdens than WT after ischemia (Fig. 8G). CLP290 10’ treatment did not improve the efficacy of PB in S940A^+/+^ mice but significantly increased 1^st^ hour seizure burden when compared to untreated S940A^+/+^ mice (Fig. 8H). During development, mice undergo a reduction in ischemic seizure susceptibility from P7 to P10 that is associated with a robust upregulation of KCC2 and S940 phosphorylation (*15, 26*). S940A^+/+^ mice did not undergo a developmental decrease in ischemic seizure susceptibility at P10 when compared to P7 S940A^+/+^ mice (Fig. 8I; P=0.57). These results suggest that both the CLP290-mediated KCC2 functional enhancement and the age-dependent reduction in ischemic seizure susceptibility are dependent upon S940 phosphorylation.

### S940A^+/+^ pups are susceptible to status epilepticus and death

To determine if the ability for KCC2 phosphorylation to modulate seizure susceptibility was specific to ischemic seizures, WT, S940A^+/+^, and T1007A^+/+^ P12 pups underwent the 3h PTZ challenge (Fig. 8J-O). As described previously (Fig. 4), pups were administered three doses of PTZ (20, 20, and 40 mg/kg) 1h apart. S940A^+/+^ mice immediately progressed into status epilepticus and died before the 2^nd^h (Fig. 8 J-O). This indicated that the prevention of S940 phosphorylation increased the risk of sudden unexpected death in epilepsy (SUDEP)-like phenomenon (*50*). In contrast, T1007A^+/+^ mice were resistant to the PTZ induced seizures when compared to WT (Fig. 8 M). Additionally, the assessment of righting reflex as a neurodevelopmental milestone identified that only S940A^+/+^ mice had a significantly impaired righting reflex at P7 but not at P12 (Fig. 8P). These data suggest that KCC2 phosphorylation controls seizure susceptibility during development.

## Materials and Methods

### Experimental Paradigm

In CLP290 and CLP257 experiments, all pups regardless of treatment group received a loading dose of PB (25mg/kg) dissolved in 100% isotonic phosphate-buffered saline (PBS) delivered via intraperitoneal (IP) injection at 1h. All injections regardless of the treatment group, drug, or age were administered using Hamilton syringes. All drugs were prepared the day of experiments. CLP290 and CLP257 were both dissolved in 45/55% 2-Hydroxypropyl-β-cyclodextrin (HPCD)/PBS with a pH range between 7.2-7.5.

To assess the efficacy of CLP290 treatment at P7, pups were assigned to the PB-only, post, primed, or pre-treatment groups (Fig. 1). PB-only treatment was defined by the single administration of PB at 1h without further intervention. Post treatment was denoted by administration of CLP290 immediately following unilateral carotid ligation with PB at 1h. 5, 10, or 20mg/kg CLP290 was administered via IP injections to P7 pups in the post treatment groups (i.e. P7 CLP290 Post 5, 10, and 20). The primed treatment group was defined by the administration of CLP290 4h preceding unilateral carotid ligation and another dose immediately following unilateral carotid ligation with PB at 1h. The pretreatment group was denoted by the single administration of CLP290 4h before unilateral carotid ligation with PB at 1h after ligation (i.e. P7 CLP290 Pre 20”).

As a carbamate prodrug of CLP257, CLP290 has an improved bioavailability in P7 CD-1 pups when compared to CLP257. To investigate the differential effects of CLP290 and CLP257 on neonatal seizure suppression, P7 pups were administered 10mg/kg CLP257 in a primed-treatment group with PB at 1h after ligation. At P10, when seizures after unilateral carotid ligation are responsive to PB, the efficacy of 10mg/kg CLP290 to improve seizure suppression was investigated. P10 pups were either assigned to the post, primed, or pretreatment groups with PB 1h after ligation.

### Animals

All experimental procedures and protocols were conducted in compliance with guidelines by the Committee on the Ethics of Animal Experiments (Permit Number: A3272-01) and were approved by the Animal Care and Use of Committee of Johns Hopkins University. CD1 litters were purchased from Charles River Laboratories (Wilmington, MA.). Newly born CD-1 litters (n=10) were delivered with a dam at postnatal day three or four and allowed to acclimate. S940A^+/+^ and T906A/T1007A^+/+^ mice were a gift from Stephen J. Moss Laboratories at the Tufts University School of Medicine. Equivalent numbers of male and female pups were introduced into the study. All mice were housed on a 12h light-dark cycle with food and water provided *ad libitum*. For surgical procedures and western blotting see Supplement.

### *In vivo* video-EEG recording and analyses

EEG recordings were acquired using Sirenia Acquisition software with synchronous video capture (Pinnacle Technology Inc. KS, USA). Data acquisition was done with sampling rates of 400Hz that had a preamplifier gain of 100 and the filters of 0.5Hz high-pass and 50Hz low-pass. The data were scored by binning EEG in 10s epochs. Similar to our previous studies(*15*), seizures were defined as electrographic ictal events that consisted of rhythmic spikes of high amplitude, diffuse peak frequency of ≥7-8Hz (i.e.; peak frequency detected by automated spectral power analysis) lasting ≥6s (i.e.; longer than half of each 10s epoch). Short duration burst activity lasting <6s (brief runs of epileptiform discharges) was not included for seizure burden calculations similar to previous studies in the model. Mean time spent seizing for 1^st^h baseline seizure burden vs. 2^nd^h post-PB seizure burden was quantified in seconds. Mean seizure suppression was calculated using Equation 1:

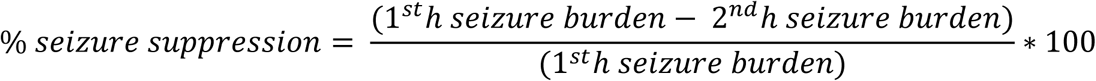

Mean ictal events and ictal durations (seconds/event) were calculated for 1^st^h vs. 2^nd^h.

## Statistics

For all experiments, the quantification and analysis of data were performed blinded to the genotype, sex, and treatment conditions. All statistical tests were performed using GraphPad Prism software. Two-way analysis of variance (ANOVA) was performed with Tukey’s post hoc correction. One-way ANOVA was performed with Dunnett’s post hoc correction. Paired and unpaired t-tests were two-tailed. Survival analysis was performed by a Mantel-Cox test. Data are represented as bar graphs representing the mean, with dot plots representing each individual data point. Errors bars are ± 1 standard error of mean. P values for ≤0.05 are reported.

## Discussion

KCC2 is one of the key regulators of intracellular chloride (*7, 51*). However, diverse mechanisms regulate KCC2 membrane insertion and Cl^−^ extrusion capacity (*11, 12*). Modulating KCC2 function to enhance inhibition has become a focused area of research to help identify therapeutic targets for pathologies with documented KCC2 degradation or hypofunction. Enhancement of KCC2 membrane stability and function could reestablish synaptic inhibition in seizing neonatal brains when positive GABAR modulators like PB fail to curb severe recurrent seizures. The goal would be to acutely rescue KCC2 function by preventing further degradation and maintaining KCC2 membrane stability. For translational applications, this strategy would target pathologies with documented KCC2 hypofunction such as recent reports of significant reduction in KCC2-positive cells in postmortem cerebral samples from preterm infants with white matter injury (*52*). This strategy is distinct from overexpression or enhancement of KCC2 function in neurons with stable endogenous KCC2 expression which could be detrimental to developing brains (*53, 54*). Moderate and severe HIE seizures are clinically associated with significant hourly/daily seizure burdens which tend to cluster into high-seizing and non-seizing periods (*4, 55*), however they are transient in nature. This clustering phenotype makes continuous EEG monitoring during the acute period a necessity and gold standard (*56, 57*) to determine both the true severity of the seizures and the efficacy of ASM interventions. An acute protocol of rescuing KCC2 hypofunction during this critical period would help mitigate both the refractory neonatal seizures and their long-term consequences.

For this reason we tested acute intervention protocols for multiple graded doses of the KCC2 functional enhancer CLP290 both as post and primed dosing in our CD-1 mouse model of refractory neonatal seizures. Our seminal results show that CLP290 is significantly efficacious in reversing PB-resistant seizures at P7 in a dose-dependent manner that plateaus at the highest dose tested in this study. The ability of CLP290 to rescue both KCC2 expression and -S940 phosphorylation at 24h post-ischemia were similar to those reported with the TrkB antagonist ANA12 but also distinct since CLP290 had no effect on subduing the ischemia-induced TrkB activation (*17*). Further its ability to significantly increase KCC2 membrane insertion in P7 brain sections and inability subdue ischemic seizures in the mutant KCC2 S940A^+/+^ pups support its role as a KCC2 functional enhancer acting through its S940 phosphorylation site.

### Novel anti-seizure protocols that help mitigate epileptogenesis

The long-term effects of refractory seizures and ASM interventions are major factors to consider for neonatal antiseizure management (*21, 22*). It has been shown that chemoconvulsant induced seizures upregulate KCC2 expression (*32, 58*). Therefore, an initial low-dose PTZ would be expected to upregulate KCC2 in P12 naïve pups likely making them resistant to additional seizures. The proof of concept data reported here in the naïve pups for this novel assay supported this reasoning. The failure of the initial sub-threshold PTZ dose to confer this resistance in the PB-only group indicated a heightened susceptibility to epileptogenesis at P12. In contrast, the P7 CLP290 treated pups showed significant protection from this epileptogenesis. Neonatal ischemia results in ∼20-40% decrease in KCC2 expression which recovers endogenously over the next few days and catches up to its developmental trajectory (*15*). The ability to rescue KCC2 hypofunction acutely during this period could help mitigate the onset of long-term impairments in circuit function opening up a promising window for transient therapy aimed at rescuing KCC2 hypofunction.

### KCC2 hypofunction can independently initiate and aggravate neonatal seizures

To elucidate the critical role of KCC2 in neonatal seizure susceptibility, the effect of the selective KCC2 inhibitor VU was tested in naïve CD-1 pups at P7. During development KCC2 undergoes a significant increase in expression (*6*) associated with a phosphorylation profile that coincides with a maturational increase in Cl-extrusion capacity (*23*). The dose-dependent emergence of spontaneous epileptiform events with graded severity in the naïve P7 CD-1 pups highlighted the significant role of KCC2 function in preventing ictogenesis in the immature brain (*59*). KCC2 antagonism during ischemic seizures further aggravated seizure severity at P7, supporting its role in the severity of seizure burdens. The immature brain has an inherent susceptibility to excitotoxic injury with propensity for seizures (*1, 59*). Our data indicate that KCC2 function in the neonatal brain plays a significant role in maintaining the balance between excitation and inhibition.

### Novel targets to rescue KCC2 hypofunction are needed

Currently there are very few studies evaluating CLP290 efficacy in models of neurological disease and there is an urgent need for the discovery of additional novel KCC2 functional enhancers. The De Koninck group originally identified CLP257 as a KCC2 functional enhancer using high throughput screening (*20*). The prodrug CLP290 was shown to have improved pharmacokinetics over CLP257 and rescued neuropathic pain in a rat model. *In vitro* studies proposed that the active drug CLP257 does not directly modulate KCC2 activity but potentiates GABA_A_R activity (*60, 61*). Regardless, CLP290 has been shown to improve outcomes in models of neuropathic pain and spinal cord injury associated with KCC2 hypofunction (*20, 62, 63*). The results of this current study highlight the significant role of KCC2 and its phosphorylation sites in neonatal seizure severity and mechanisms underlying CLP290 ASM efficacy using both *in vivo* and *in vitro* protocols. The identification of newer and more efficient KCC2 functional enhancers targeting these phosphorylation sites are needed.

The mechanisms by which CLP290 enhances KCC2 function in neurons are not well understood (*20, 60, 61*). We investigated the role of the known phosphorylation sites on KCC2 those that either positively or negatively regulate KCC2 membrane stability and Cl^−^ extrusion capacity, in the two mutant mice. The S940A^+/+^ pups not only showed an increase in seizure susceptibility to the P7 ischemic insult, but importantly we documented the occurrence of spontaneous epileptiform discharges in the naïve mutant pups at P7. S940A^+/+^ pups did not show the age-dependent (P7 vs. P10) differences in ischemic seizure susceptibility highlighting the developmental role of S940 phosphorylation in hyperexcitability of the immature brain. The significance of this finding is highlighted in recent reports of KCC2 mutations in early-onset epileptic syndromes (*9*), in which mutant KCC2 is associated with a reduction in S940 phosphorylation (*46*) and impaired Cl^−^ extrusion (*47, 64*). In contrast, T1007A^+/+^ mice showed a resistance to post-ischemic seizure susceptibility as compared to their WT littermates following P7 ischemic insults.

Additionally, when the repeated-dose PTZ protocol was tested in both mutant strains at P12, the high seizure burdens and early mortality detected in S940A^+/+^pups and low seizure susceptibility of the T1007A^+/+^ pups may further support the role of the two sites in the evolution of epileptogenesis following P7 neonatal seizures in the CD-1 model.

### Activation of KCC2 homeostatic compensatory mechanisms with bolus doses of CLP290

In contrast to the graded ASM responses of CLP290 for the lower doses tested, the anti-seizure responses identified for the highest CLP290 dose (20mg/kg; both Pre and Post) tested in this study showed differences. First hour baseline seizure burdens which remained similar to ligate-only group for the lower doses of CLP290 were significantly suppressed with the post-ligation bolus dose of 20mg/kg (Suppl Fig 1). This alteration of the baseline seizure susceptibility at P7 with 20mg/kg of CLP290 took away the graded nature on the reversal of PB-refractoriness seen with the 5 vs. 10mg/kg doses (Fig.1).

KCC2 can autoregulate its Cl^−^ extrusion capacity using positive and negative modulators that activate its multiple phosphorylation sites (*23*). It is also known that modest KCC2 hypofunction can play a significant role in the emergence of neurological disorders (*37*). In contrast, overexpression of KCC2 in neurons could not only be detrimental but also trigger endogenous homeostatic pathways for the negative regulation of KCC2 function. Interestingly, pushing KCC2 function beyond its physiological levels was found to be difficult in simulation studies (*11*). The high bolus dose of CLP290 tested here was found to activate one such endogenous mechanism by which KCC2 upregulation as denoted by a significant change in ischemic seizure susceptibility at P7 was homeostatically countered by a significant upregulation of T1007 phosphorylation at 24h. The upregulation of T1007 in the CLP290 20mg/kg treatment group was not detected in the untreated P7 CD-1 ischemia group nor the with the lower doses of CLP290 and therefore may indicate a homeostatic regulation of KCC2 function induced by CLP290 high bolus dose at 24h that was independent of the initial ischemic insult. The findings of the CLP257-mediated T007 phosphorylation *in vitro* versus *in vivo* experiments indicate a temporal regulation of the homeostatic response with T1007 upregulation which needs further investigation.

### Novel insights for the next generation of the next generation of KCC2 enhancers

Recent studies have shown that during induced status epilepticus, reduced Cl^−^ extrusion capacity and exacerbated activity-dependent Cl^−^ loading can result in GABAergic transmission being ictogenic (*14*). Optogenetic stimulation of GABAergic interneurons in this status epilepticus-like state enhanced the epileptiform activity in a GABA_A_R dependent manner (*65*) indicating GABA-mediated depolarization. These findings support data both from our model and clinical reports where the when initial loading-dose of PB fails to curb seizures, additional PB doses do not help rescue the refractoriness (*2, 66*). Additionally, given the toxicity of high-dose PB on neonatal brains such protocols may be counterproductive in the short and long-term (*67*–*69*).

KCC2 hypofunction is emerging as a significant cause underlying impaired inhibition in multiple neurological disorders (*5, 52, 65*). Interestingly, the research into both refractory seizures and refractory spinal nerve pain has identified KCC2 hypofunction as a common underlying cause. Here we have shown that enhancing KCC2 function in a well-characterized preclinical model of refractory seizures can rescue not only the acute PB-refractoriness but also help mitigate epileptogenesis with early intervention. Although additional studies are needed to investigate the direct and indirect effects of enhancing Cl^−^ extrusion capacity of KCC2 in the immature brain, the novel findings reported here highlight the role of KCC2 hypofunction and its phosphorylation sites in HIE-related refractory seizures and an evidence based focus on targeting KCC2 function in development of future translational strategies.

## Acknowledgements

We thank Professor Stephen J. Moss (Tufts University School of Medicine) for the generous transfer of the S940A^+/+^ and T1007A^+/+^ mice (MTA). We thank Dr. Tarek Deeb (Tufts University School of Medicine) and Dr. Yvonne Moore (Tufts University School of Medicine) for useful discussions and technical advice. We thank Professor Yves De Koninck (Université Laval) for generously sharing aliquots of CLP290 (MTA). We thank Dr. Rana Rais from the Johns Hopkins Drug Discovery Core for the HPLC experiments.

## Author Contributions

SDK conceived the project. BJS, PAK, BMC, and SDK acquired data. BJS, PAK, and SDK analyzed data. BJS, PAK, and SDK wrote the paper. All authors have seen and approved the manuscript, and this manuscript has not been published elsewhere.

## Supplementary Materials for

## Materials and Methods

### Unilateral carotid ligation

A comprehensive protocol for unilateral carotid ligation and neonatal video-EEG recordings has been published (*71*) At P7 or P10, animals were subjected to permanent unilateral ligation (without transection) of the right common carotid artery using 6-0 surgisilk (Fine Science Tools, BC Canada) under isoflurane anesthesia. The outer skin was closed with 6-0 monofilament nylon (Covidien, MA), and lidocaine was applied as local anesthetic. Animals were implanted with 3 subdermal EEG scalp electrodes: 1 recording and 1 reference overlying the bilateral parietal cortices, and 1 ground electrode overlying the rostrum. Wire electrodes (IVES EEG; Model # SWE-L25 –MA, IVES EEG solutions, USA) were implanted subdermally and fixed in position with cyanoacrylate adhesive (KrazyGlue). Pups recovered from anesthesia over a few minutes. Animals were tethered to a preamplifier within a recording chamber for 2h of continuous vEEG recording and were maintained at 36°C with heated isothermal pads. At the end of the recording session, sub-dermal electrodes were removed, and the pups were returned to the dam.

### CLP290 plasma and brain availability in neonatal mice

Standards for HPLC were created using CLP257 (MilliporeSigma, USA) and CLP290 (Yves De Koninck Lab). P7 and P10 naïve pups of both sexes were administered CLP290 IP as three treatment groups: 10mg/kg, 20mg/kg, or vehicle. After 4h pups were anesthetized with chloral hydrate (90 mg/ml; IP), and transcardiac blood samples (100μL) were collected. The same pups were transcardially perfused with ice cold PBS, and the whole fresh brains harvested. Brain samples were flash frozen in dry ice, homogenized using a sonicator, and stored at −80°C. Blood and brain samples from CLP290 treated and naïve pups were analyzed for CLP290 and CLP257 concentrations via HPLC using a C18 column and 10/90 organic/aqueous mobile phase.

### VU 0463271

To assess the role of KCC2 inhibition in neonatal seizure severity at P7 after ligation, the potent and selective KCC2 inhibitor VU 0463271 (VU) was administered at 1h (0.5mg/kg; IP) in lieu of PB (Figure 1). VU was dissolved in 20/80% dimethyl sulfoxide (DMSO)/PBS solution. To assess to role of KCC2 antagonism in neonatal seizure occurrence, naïve P7 or P10 pups underwent vEEG with 0.25mg/kg VU administered at the start of recording in lieu of ligation and 0.5mg/kg VU administered at 1h in lieu of PB.

### P12 PTZ Challenge

To investigate the long-term effects of CLP290 treatment, P12 pups underwent a three hour vEEG recording during a pentylenetetrazole (PTZ; dissolved in 100% PBS) challenge. At P7 pups were either naive, PB-only, or P7 CLP290 10’. These P7 pups then underwent a PTZ challenge at P12. All pups were administered PTZ at the beginning of the vEEG recording (20mg/kg; IP), at 1h (20mg/kg; IP), and at 2h (40mg/kg; IP).

### Western blot analysis at 24h post-ligation

All animals for immunochemical characterizations were anesthetized with chloral hydrate (90 mg/ml; IP) before being transcardially perfused with ice cold saline. The whole fresh brains were removed, the cerebellum was discarded, and the left and right hemispheres were separated. Brains were stored at −80°C in preparation for further processing. Brain tissue homogenates were made and suspended in TPER cell lysis buffer containing 10% protease/phosphatase inhibitor cocktail. Total protein amounts were measured using the Bradford protein assay (Bio-Rad, Hercules, CA, USA) at 570nm and the samples diluted for 50μg of protein in each sample. 20μL of protein samples were run on 4-20% gradient tris-glycine gels (Invitrogen, Gand Island, NY, USA) for 120min at 130V and were transferred onto nitrocellulose membranes overnight at 20V. After the transfer, the nitrocellulose membranes underwent a 1h blocking step in Rockland buffer before 6h incubation with primary antibodies (for all antibody RRIDS, see Key Resources Table): mouse α-KCC2 (1:1000, Millipore), rabbit α-phospho-KCC2-S940 (1:1000 Aviva Systems Biology), rabbit α-phospho-KCC2-T1007 (1:1000; Phospho solutions) mouse α-TrkB (1:1000, BD Biosciences), rabbit α-phospho-TrkB-Y816 (1:500, Millipore), and mouse α-actin (1:10000, LI-COR Biosciences). Membranes were then incubated with fluorescent secondary antibodies (1:5000, goat α-rabbit and goat α-mouse, Li-Cor Biosciences, USA). Chemiluminescent protein bands were analyzed using the Odyssey infrared imaging system 2.1 (LI-COR Biosciences). The optical density of each protein sample was normalized to their corresponding actin bands run on each lane for internal control. Mean normalized protein expression levels were then calculated for respective left and right hemispheres. The expression levels of the proteins of interest in ipsilateral hemispheres were normalized to the same in contralateral hemispheres for each pup to examine hemispheric percent change of protein expression.

### Surface protein-seperation

1mm coronal brain slices were obtained from P7 CD-1 mice and recovered for 45min at 34°C with oxygenation (95%/5% O_2_/CO_2_). After recovery, slices were incubated with CLP290 or CLP257 at 34°C with oxygenation for 40min. Slices were placed in TPER cell lysing buffer with HALT protease and phosphatase inhibitors and homogenized via sonication. After 30min incubation on ice, protein lysates were ultracentrifuged at 210,000xg (TLA-120.2 rotor, Beckman Coulter Life Sciences), and supernatants were collected as the cytosolic components. Pellets were resuspended in TPER/HALT buffer and ultracentrifuged; supernatant was discarded as wash fraction. Pellets were resuspended in TPER/HALT buffer and collected as the membrane components. Membrane and cytosolic components underwent Bradford analysis and Western blotting for protein quantification. Plasma membrane proteins were normalized to TfR. Cytosolic proteins were normalized to β-actin.

## Abbreviations

PB: Phenobarbital
ASM: Antiseizure medication
EEG: Electroencephalography
KCC2: K-Cl co-transporter 2
NKCC1: Na-K-Cl cotransporter 1
GABAR: GABA receptor
HIE: hypoxic-ischemic encephalopathy
TrkB: Tropomyosin receptor kinase B

---

**Supplemental Fig. 1.**
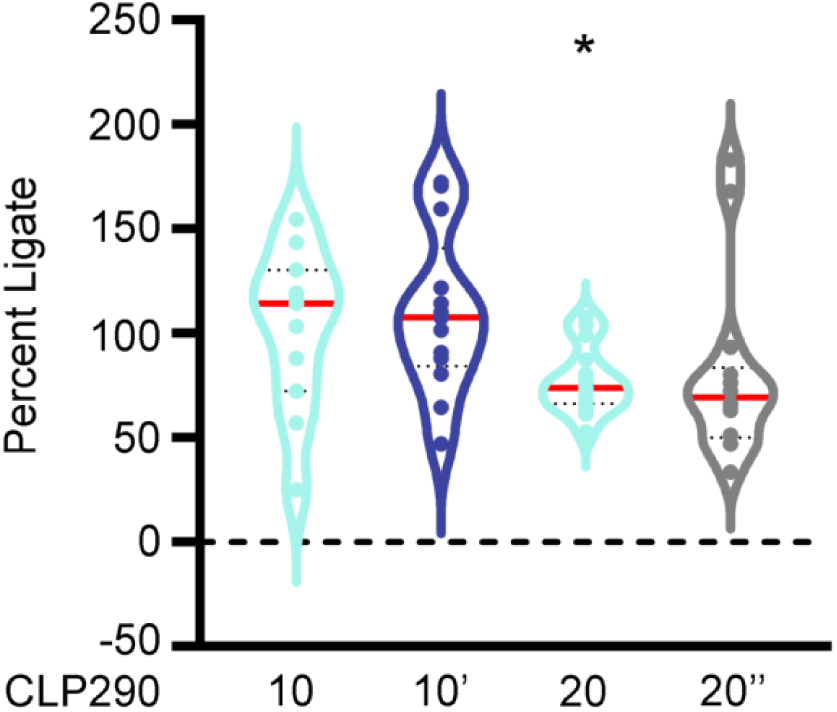
CLP290 20 Post reduced first hour seizure burden. 1^st^ hour seizure burdens for 10,10’,20, and 20” doses of CLP290. Violin plots show all data points as percent PB-only 1^st^ hour seizure burden. P=0.0477, two-tailed *t*-test vs. PB-only 1^st^ hour seizure burden.

**Supplemental Fig. 2.**
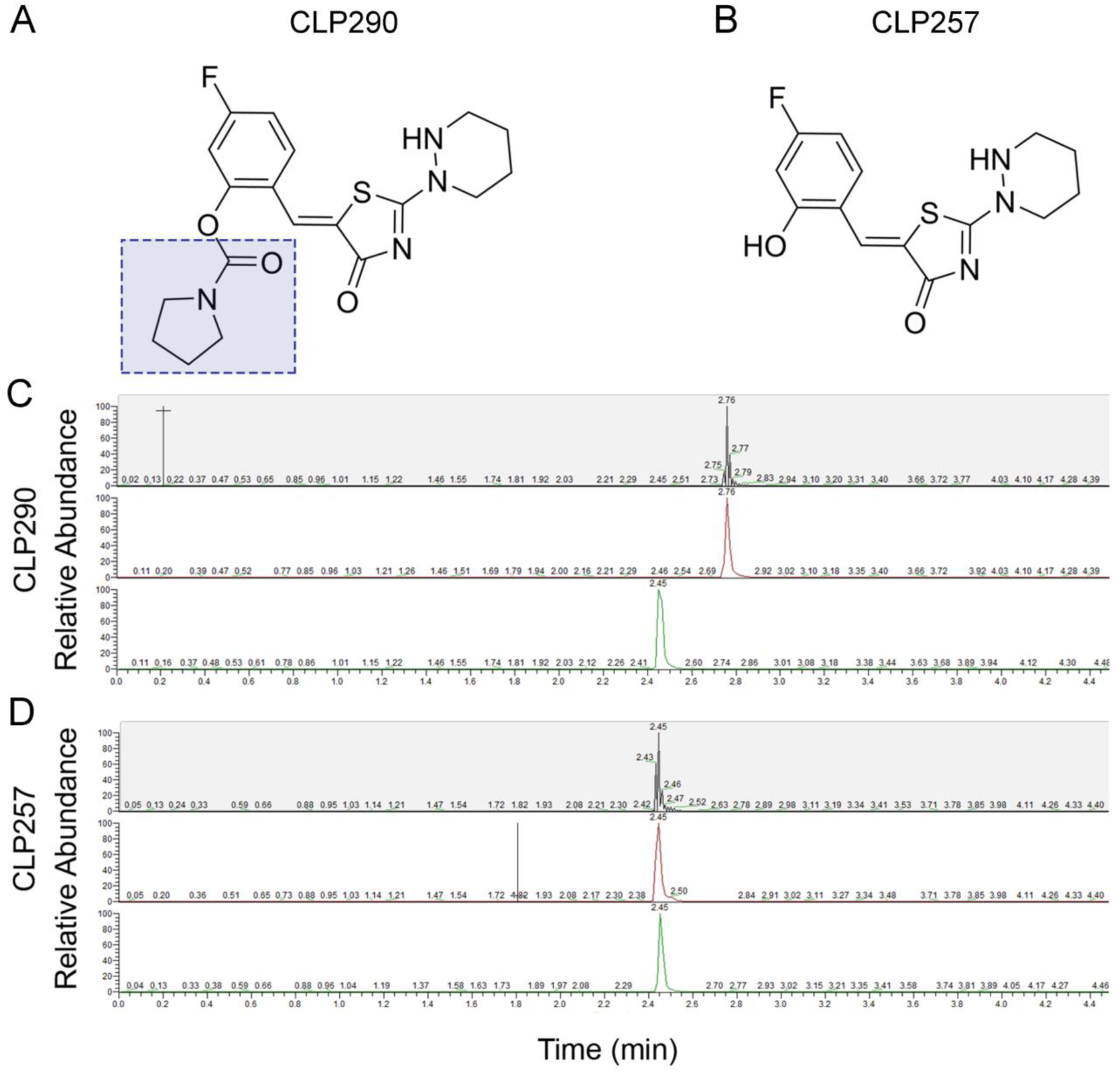
Characteristic peaks of CLP290 and CLP257 on HPLC. (**A**) CLP290 is the carbamate prodrug of (**B**) CLP257. (**C**) The characteristic peaks of CLP290 (Pharmablock) and (**D**) CLP257 (Sigma) on HPLC.

**Supplemental Fig. 3.**
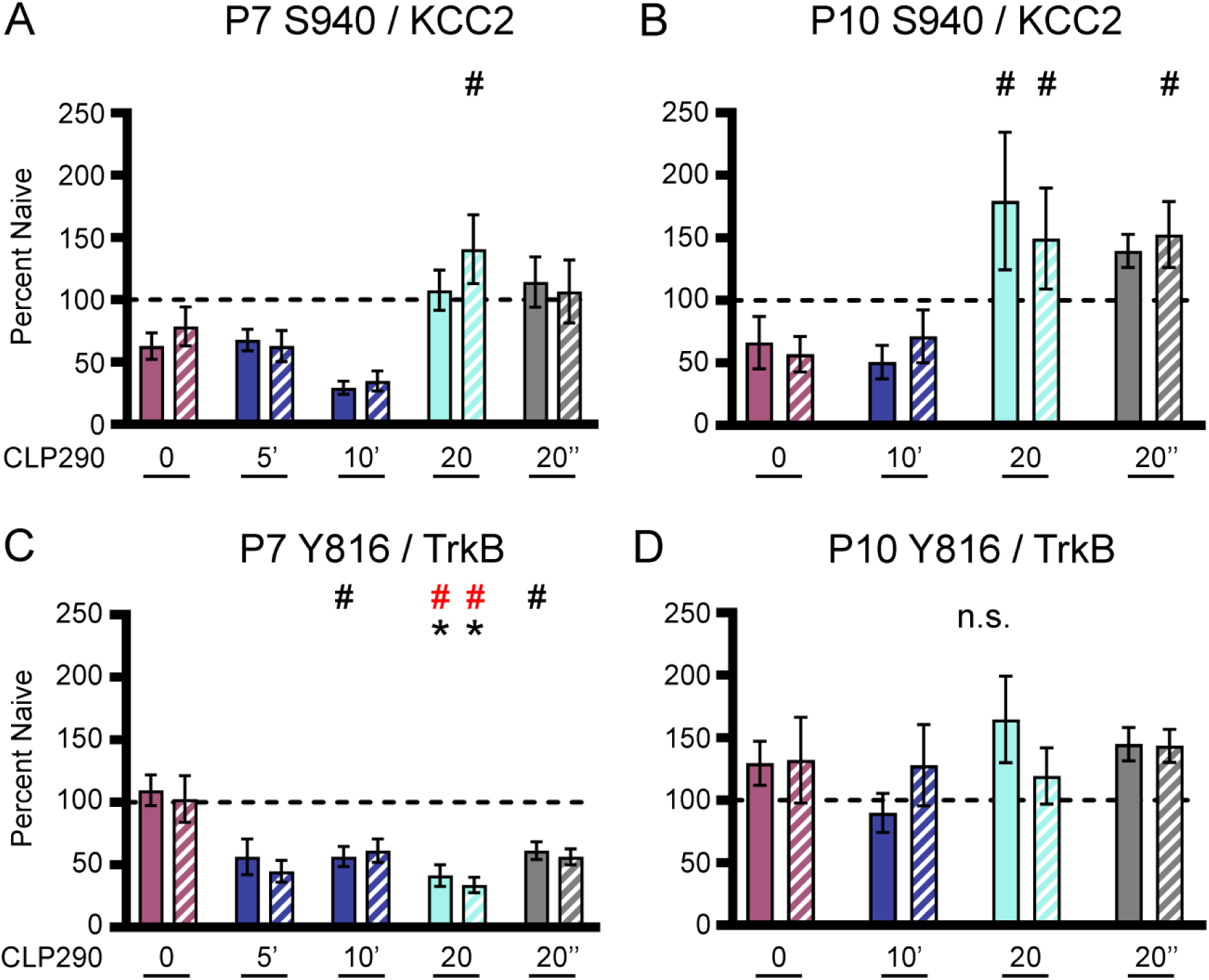
CLP290 reduced Y816 activation. (**A**) S940/KCC2 ratios 24h after P7 ischemic neonatal seizures plotted as left and right hemispheres. (**B**) S940/KCC2 ratios 24h after P10 ischemic neonatal seizures plotted as left and right hemispheres. (**C**) Y816/TrkB ratios 24h after P7 ischemic neonatal seizures plotted as left and right hemispheres. (**D**) Y816/TrkB ratios 24h after P10 ischemic neonatal seizures plotted as left and right hemispheres. *P<0.05 and *P<0.001 by 1-way ANOVA vs. Naive. #P<0.05 and #P<0.001 vs. PB-only P7 pups: Naïve n=27; PB-only n=18; 5 Post n=3; 10 Post n=9; 20 Post n=13; 5 Primed n=4; 10 Primed n=11; 20 Pre n=9. P10 pups: Naïve n=18, PB-only n=11, 10 Post n=6, 20 Post n=6, 10 Primed n=5, 20 Pre n=7.

**Supplemental Fig. 4.**
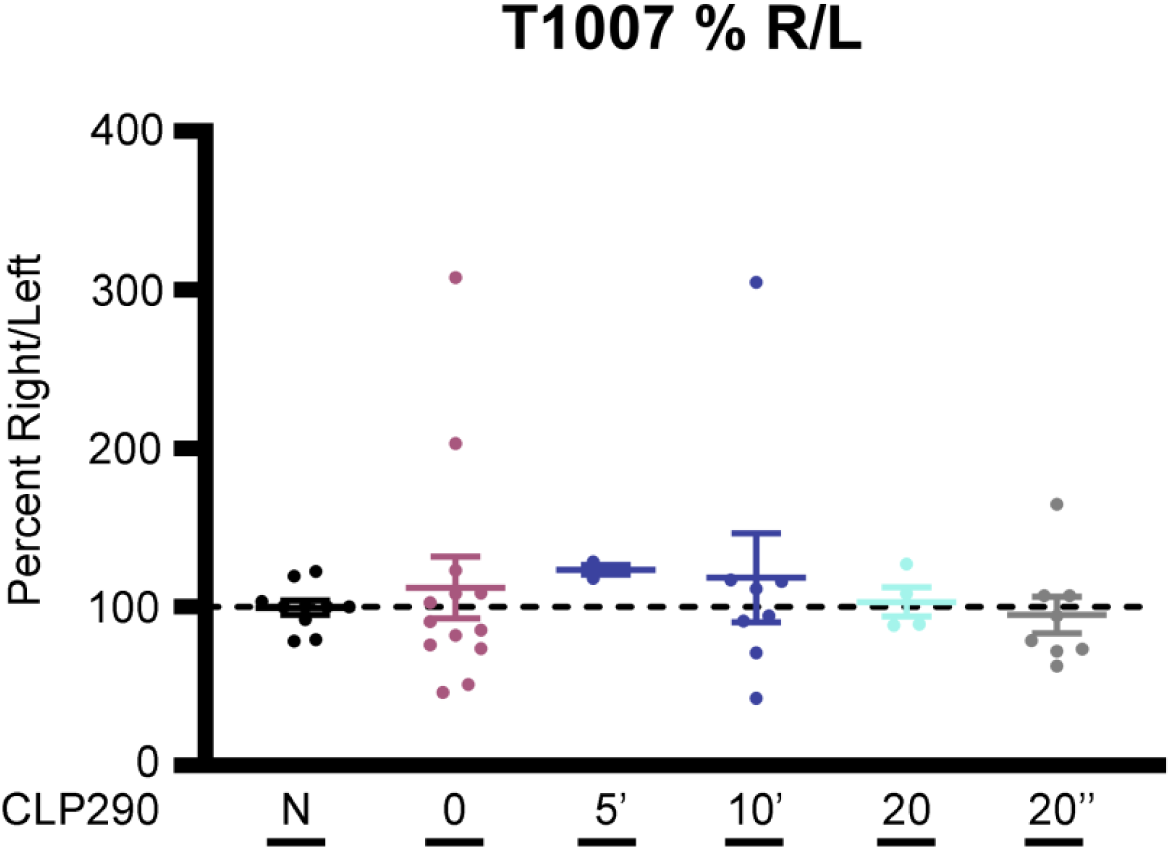
T1007 is not activated by ischemia. T1007 expression as percent ipsilateral/contralateral plotted as percent naïve.

| Reagent type (species) or resource | Designation | Source Reference | Identifiers | Additional Information |
| --- | --- | --- | --- | --- |
| Genetic reagent (M. musculus) | CD-1 | Charles River | 022 | N/A |
| Genetic reagent (M. musculus) | S940A | Silayeva et al. 2015 | N/A | Dr. Stephen Moss, Tufts University School of Medicine |
| Genetic reagent (M. musculus) | T1007A | Moore et al. 2018 | N/A | Dr. Stephen Moss, Tufts University School of Medicine |
| Chemical compound, drug | Phenobarbital (PB) | MilliporeSigma | P5178 | N/A |
| Chemical compound, drug | CLP290 | Gagnon et al. 2013 | N/A | Dr. Yves De Koninck, Université Laval |
| Chemical compound, drug | CLP290 | PharmaBlock | PBSQ8214 | N/A |
| Chemical compound, drug | CLP257 | MilliporeSigma | SML1368 | N/A |
| Chemical compound, drug | Pentylene-tetrazol (PTZ) | MilliporeSigma | P6500 | N/A |
| Chemical compound, drug | VU0463271 (VU) | Tocris | 4719 | N/A |
| Chemical compound, drug | DMSO | Sigma | D8418 | N/A |
| Chemical compound, drug | HPCD | Sigma | H107 | N/A |
| Software, algorithm | Graphpad Prism | Graphphad Software | RRID:SCR_002798 | 8 |
| Software, algorithm | Sirenia | Pinnacle Technology | pinnaclet.com/sirenia | 3-Channel EEG/EMG Tethered Mouse System |
| Antibody | mouse $\alpha$ KCC2 | Aviva Systems Biology OASE00240 | AB_2721238 | 1:1000; WB |
| Antibody | rabbit $\alpha$ pKCC2-S940 | Aviva Systems Biology OAPC00188 | AB_2721198 | 1:1000; WB |
| Antibody | rabbit $\alpha$ pKCC2-1007 | PhosphoSolutions p1551-1007 | AB_2716769 | 1:1000; WB |
| Antibody | mouse $\alpha$ TrkB | BD Biosciences 610102 | AB_397508 | 1:1000; WB |
| Antibody | rabbit $\alpha$ pTrkB-Y816 | Millipore ABN1381 | AB_2721199 | 1:500; WB |
| Antibody | mouse $\alpha$ actin | LI-COR Biosciences 926-42213 | AB_2637092 | 1:10000; WB |
| Antibody | mouse $\alpha$ Transferrin Receptor | ThermoFisher Scientific | AB_2533029 | 1:500; WB |
| Antibody | goat $\alpha$ mouse IgG, IRDye® 800CW Conjugated | LI-COR Biosciences 926-32210 | AB_621842 | 1:5000; WB |
| Antibody | goat $\alpha$ rabbit IgG Antibody, IRDye® 680LT Conjugated | LI-COR Biosciences 926-68021 | AB_10706309 | 1:5000; WB |

